# OnePetri: accelerating common bacteriophage Petri dish assays with computer vision

**DOI:** 10.1101/2021.09.27.460959

**Authors:** Michael Shamash, Corinne F. Maurice

## Abstract

**Introduction:** Bacteriophage plaque enumeration is a critical step in a wide array of protocols. The current gold standard for plaque enumeration on Petri dishes is through manual counting. This approach is time-intensive, has low-throughput, is limited to Petri dishes which have a countable number of plaques, and can have variable results upon recount due to human error.

**Methods:** We present OnePetri, a collection of trained machine learning models and open-source mobile application for the rapid enumeration of bacteriophage plaques on circular Petri dishes.

**Results:** When compared against the current gold standard of manual counting, OnePetri was significantly faster, with minimal error. Compared against two other similar tools, Plaque Size Tool and CFU.AI, OnePetri had higher plaque recall and reduced detection times on most test images.

**Conclusions:** The OnePetri application can rapidly enumerate phage plaques on circular Petri dishes with high precision and recall.

## Introduction

Bacteriophage (phage) enumeration through plaque counting is central to many assays and experiments. The classical double agar overlay plaque assay protocol has long been used to yield visible phage plaques on a solid lawn of susceptible host bacteria [1], where plaque formation is usually a direct result of bacterial infection and cell death [2]. To date, manual counting of plaques has remained the gold standard for phage enumeration and titers. In addition to being time-intensive, this method is often limited to plates with a countable number of plaques, typically less than 300, and results can vary upon recount by different individuals. To this end, several image processing techniques for automating plaque counts have been created in recent years, many of which rely on contour or edge detection to identify plaques [3–6]. Some of these tools require specific types of images, such as those obtained through fluorescence microscopy, to obtain plaque counts, increasing experimental complexity for the benefit of automation. Despite being designed to automate plaque counts, these tools often require user intervention and fine-tuning of image processing parameters to improve detection results and avoid false positives. Furthermore, most tools created for this purpose are designed to run on a desktop computer, breaking the workflow where counts would be followed by calculations and immediate experimental continuation. We thus developed OnePetri, a mobile application using a collection of trained machine learning object detection models for the rapid enumeration of phage plaques on circular Petri dishes. Using images provided by the Howard Hughes Medical Institute’s (HHMI) Science Education Alliance-Phage Hunters Advancing Genomics and Evolutionary Science (SEA-PHAGES) program [7], we successfully trained Petri dish and phage plaque object detection models which have high recall and precision. Using machine learning and computer vision, we were able to build a flexible solution which can detect diverse plaque morphologies on different types of agar media, regardless of lighting conditions, without requiring any special image capture devices or fluorescent labeling. When benchmarked against two other similar tools, CFU.AI (Apple App Store) and Plaque Size Tool [5], OnePetri was significantly faster and more accurate, reproducibly detecting hundreds of overlapping and non-overlapping phage plaques within a few seconds.

## Methods

### Image dataset curation, annotation, and preprocessing

Over 12,000 images were generously provided by the HHMI SEA-PHAGES program from the PhagesDB database [7] for use in training machine learning models. After manual curation to remove images smaller than 1,024 × 1,024 pixels in size, as the plaques in these images would be too low resolution for model training, approximately 10,000 images remained. A random subset of 98 images (75 training + 23 validation) and 38 images (29 training + 9 validation) were used when preparing the Petri dish detection and plaque detection models, respectively. Most Petri dish images had more than one Petri dish, while most plaque assay images had at least 100 plaques. Images were manually annotated (ground-truth bounding boxes drawn) using the Roboflow online platform (Roboflow Inc., Des Moines, IA, USA). Prior to export for training, annotated images for the Petri dish detection model were preprocessed to fit within 1,024 × 1,024 pixels (maintaining aspect ratio), with no additional augmentations. Annotated images for the plaque detection model were automatically preprocessed as follows: tiled into 5 rows and 5 columns, tiles resized to fit within 416 × 416 pixels (maintaining aspect ratio). The following augmentations were applied to the plaque detection training dataset, with a total of 3 outputs being produced per training example (tile): grayscale applied to 35% of images, hue shift between −45° and +45°, blur up to 2 pixels, mosaic. After augmentations (training set only) and tiling (training + validation sets), the total number of tiles used for the plaque detection model training and validation were 2,175 and 225, respectively.

### Machine learning model training and validation

The trained PyTorch models were generated using the annotated, preprocessed, and augmented dataset and the Ultralytics YOLOv5 training script (Ultralytics, Los Angeles, CA, USA) [8, 9]. When working with object detection machine learning models, the training and inference processes are generally faster and uses less video random access memory (VRAM) on the graphics processing unit (GPU) when the training images are of lower resolution. While it would be ideal to train a model using images with their native multi-megapixel resolution, this is not usually feasible given the VRAM limits on many GPUs. The following parameters were used to train the Petri dish detection model: 320 × 320 pixel image resolution (scale to fit each Petri dish image), 500 epochs, batch size of 16, YOLOv5s model (yolov5s.pt weights file), cache images enabled, default hyperparameters (hyp.scratch.yaml file). The following parameters were used to train the plaque detection model: 416 × 416 pixel image resolution (scale to fit each tile), 500 epochs, batch size of 128, YOLOv5s model (yolov5s.pt weights file), cache images enabled, default hyperparameters (hyp.scratch.yaml file).

The generated “best.pt” weights files for the trained YOLOv5 models were converted to the Apple Core ML “mlmodel” file format using the coremltools Python package (version 4.1) in conjunction with a custom script provided by Hendrik Kueck (Pocket Pixels Inc., Vancouver, BC, Canada) which is available at the following GitHub repository link: https://github.com/pocketpixels/yolov5/blob/better_coreml_export/models/coreml_export.py.

### iOS mobile application development and benchmarking

The mobile application for iOS was developed using the Swift programming language (version 5) using the Xcode 12.5.1 (build 12E507) IDE (Apple Inc., Cupertino, CA, USA) with a target SDK of iOS 13. Benchmarking of the application was carried out on the iPhone 12 mini simulator available within Xcode 12 (iOS 14.5, build 18E182), as well as on a physical iPhone 12 Pro running the same operating system build as the simulator. The iOS development simulator was running on a 2020 MacBook Air with M1 chip and 16GB RAM, and which was connected to the power adapter.

### Benchmarking OnePetri

Fifty images were randomly selected from the original unprocessed dataset provided by the SEA-PHAGES team to be used in the benchmarking analysis. These images were not included in any of the model training or validation datasets for either the petri dish or plaque YOLOv5 models, meaning this is the first time the models encounter these images for inference. Images were processed sequentially in OnePetri (v1.0.1-8) and compared to the manual counting gold standard and to two other software programs. These include CFU.AI (v1.4), a free mobile application on iOS and Android originally developed in 2019 to count bacterial colony forming units (CFU), as well as Plaque Size Tool (PST), a recently published Python tool for desktop computers which detects plaques and measures their size [5]. Various methods were used to determine the time to get a result, depending on the tool: OnePetri uses code embedded within the compiled application to report runtime statistics to the debug console when running in debug mode on-device and in simulator. PST runtime was measured using the “time” command from the command line. CFU.AI runtime was approximated with a stopwatch, as the source code is not publicly available and there is no way to natively measure application runtime on iOS. All statistical analyses and data visualizations were performed using R version 4.1.0 (2021-05-18, aarch64).

### Code and data availability

The Swift source code and Xcode project for OnePetri for iOS is available under the GNU General Public License v3.0 (GPL-3.0) at the following link: https://github.com/mshamash/OnePetri. The trained machine learning models (PyTorch and Apple MLModel formats) are available at the following link: https://github.com/mshamash/onepetri-models. The training data used for the initial versions of the Petri dish and plaque detection models are available under the Attribution-NonCommercial-ShareAlike 4.0 International (CC BY-NC-SA 4.0) license on the Roboflow Universe platform: https://universe.roboflow.com/onepetri/onepetri. The benchmarking dataset, analysis scripts, and raw data are available at the following link: https://github.com/mshamash/onepetri-benchmark.

## Results

### Trained Petri dish and bacteriophage plaque object detection models are accurate and precise

Due to a lack of publicly available trained object detection models able to identify Petri dishes and phage plaques, we first set out to train two such models. Using the YOLOv5 training scripts, we developed a model to detect common circular Petri dishes in a lab environment. The training and validation datasets were comprised of 75 and 23 images, respectively. The models trained using these images performed well and as such no additional images were added to the training datasets. After 500 epochs, the model achieved 96% precision (**Figure 1A**) and 100% recall (**Figure 1B**).

**Figure 1.**
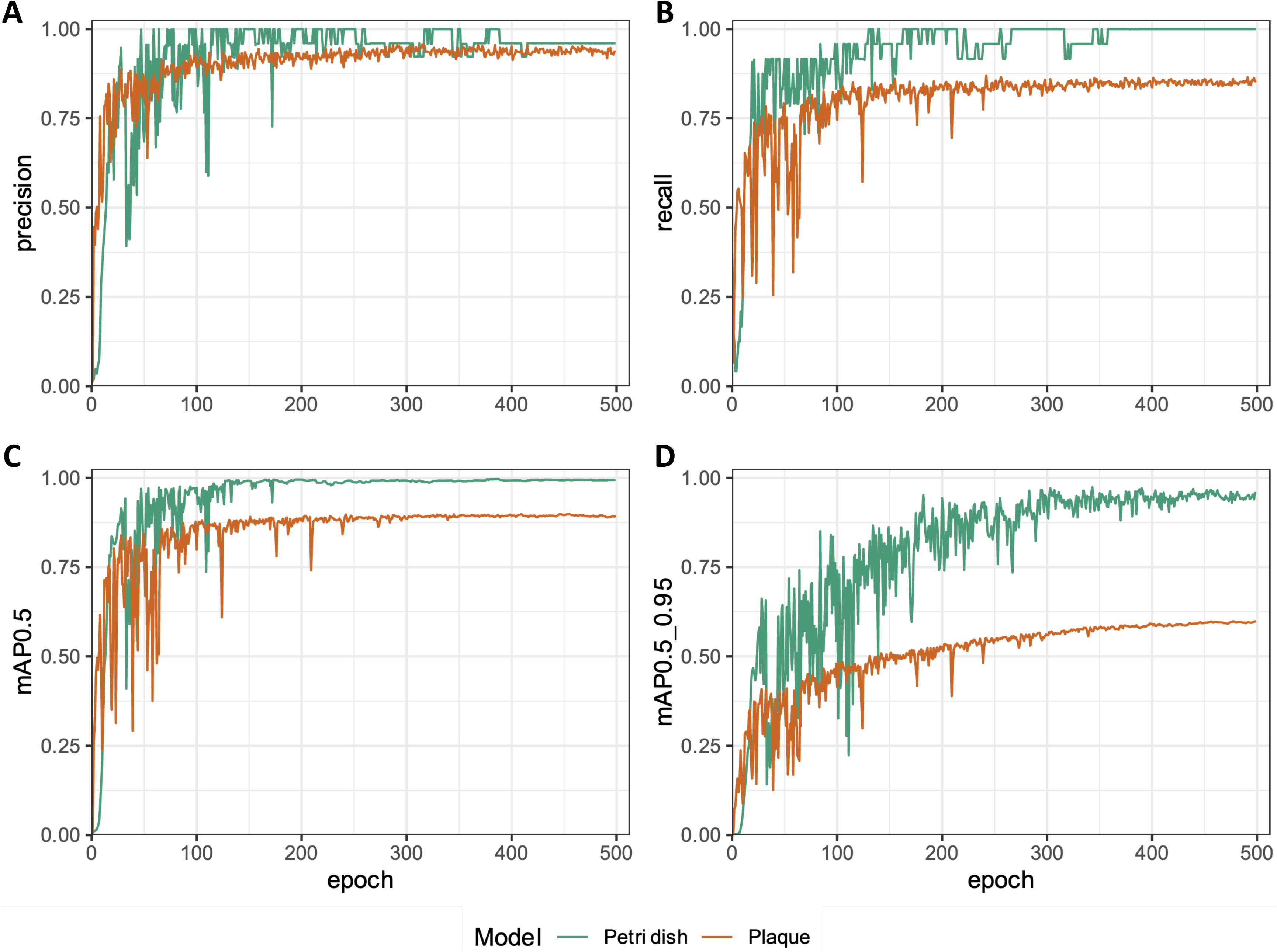
The trained Petri dish and plaque object detection models have high recall and precision after 500 training epochs. Petri dish and plaque object detection model performance metrics were recorded throughout model training using the validation datasets to test the models after each round of training, over the 500 training epochs. The following metrics were recorded and included in the figure above: **(a)** precision, **(b)** recall, **(c)** mean average precision for an intersection over union of 0.50 (mAP0.5), and **(d)** mean average precision for intersection over union ranging from 0.50 to 0.95 (step size 0.05; mAP0.5_0.95).

In computer vision, the intersection over union (IoU), also known as the Jaccard index, is a measure to evaluate how well the detected object boundaries (obtained from testing the trained model) overlap with the actual object boundaries specified prior to training [10]. The mAP@[0.5:0.95] (range of IoUs from 0.50 to 0.95, step size 0.05) and mAP@0.5 metrics are a measure of the model’s mean average precision (mAP) at the indicated IoU threshold (or range of thresholds), where detections below the threshold are not counted. Models with high mAP@[0.5:0.95] values (typically greater than 50%) are thus preferred, as this would suggest the models’ precision remains high despite increasingly stringent cutoff values for what can be considered a true positive detection. The mAP at an IoU of 0.50 (mAP@0.5) was 99.5% (**Figure 1C**), while the mAP@[0.5:0.95], was 95.7% (**Figure 1D**).

Next, to be able to detect a wide variety of plaque morphologies on diverse agar colors, we developed a model to detect phage plaques using our tiled and augmented initial dataset. The training and validation datasets were comprised of 2,175 and 225 tiles, respectively. After 500 epochs, the model achieved 93.6% precision (**Figure 1A**) and 85.6% recall (**Figure 1B**). The mAP@0.5 was 89.5% (**Figure 1C**), while the mAP@[0.5:0.95] was 59.9% (**Figure 1D**).

All performance metrics for both models plateaued after around 300 epochs, indicating that it may have been sufficient to stop training at this point.

### An image processing pipeline for rapid Petri dish detection and bacteriophage plaque enumeration

We then set out to create a mobile application wrapper for the trained models above, to allow for rapid phage plaque enumeration and assay calculations with a user-friendly interface while in the laboratory environment, without having to transfer images from a mobile phone or camera to a computer. The OnePetri mobile application was developed to fulfill this purpose and is currently available for download on the Apple App Store for free. Briefly, upon launching OnePetri, the user is first prompted to select an image for analysis (from photo library or to be taken with camera). The Petri dishes are then identified, and the user selects the Petri dish of interest to proceed with plaque enumeration analysis. This approach allows users to serially analyze multiple Petri dishes from a single image, increasing throughput. The image is cropped to the Petri dish boundaries and tiled into overlapping tiles of 416 × 416 pixels in size, where plaques are then identified serially on each tile. Finally, plaque deduplication occurs to account for plaques which may have been identified twice on overlapping tiles, and the final counts are returned to the user (**Figure 2**). Additionally, OnePetri for iOS can automatically perform the necessary calculations to obtain phage titre from Petri dishes of multiple phage dilutions as needed by the user.

**Figure 2.**
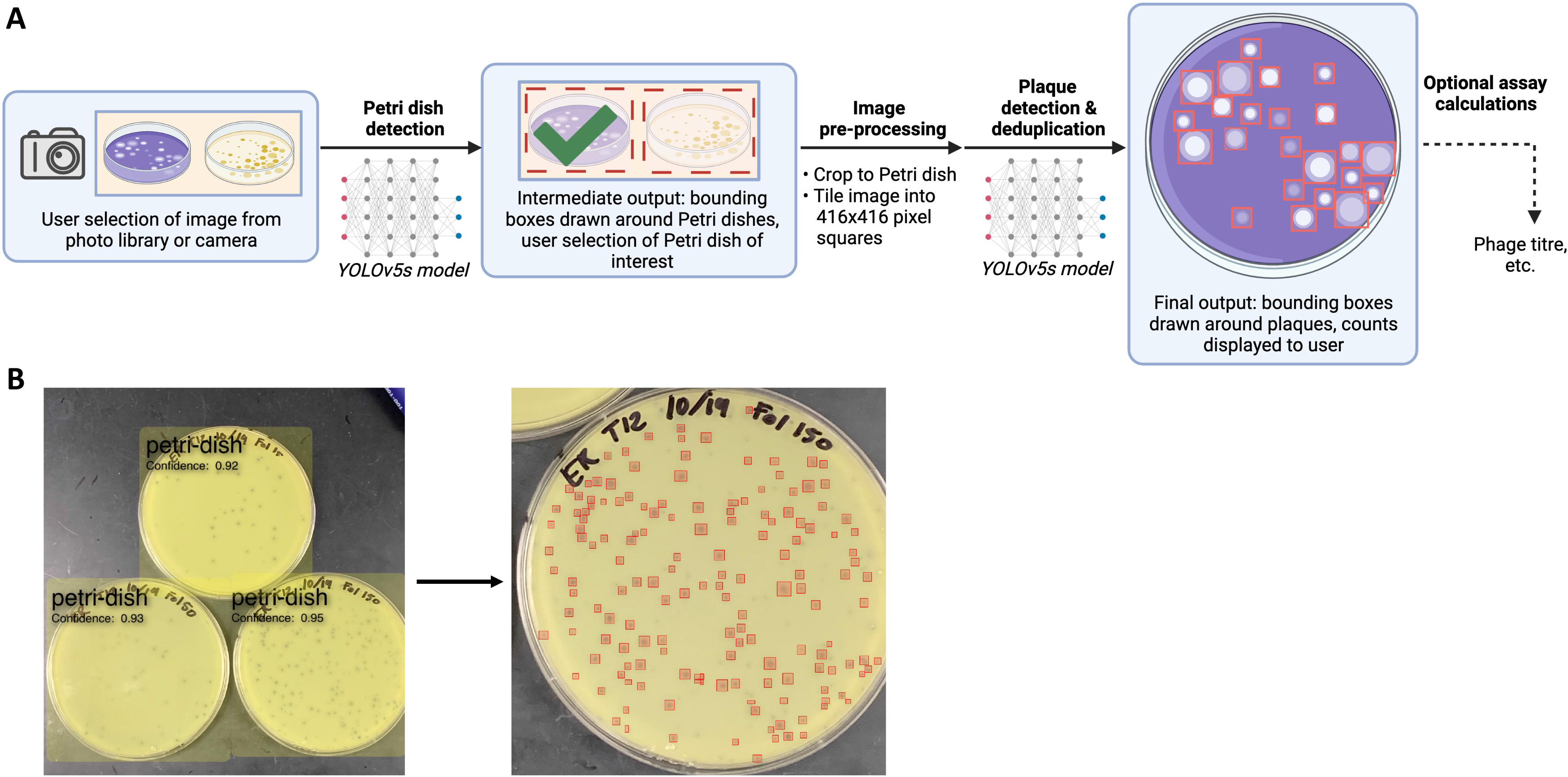
Overview of the OnePetri mobile application image processing pipeline for Petri dish detection and plaque enumeration. **(a)** Upon selecting an image for analysis, all circular Petri dishes are detected using the trained Petri dish detection model. The user selects the Petri dish they wish to analyze, and the image is cropped to fit that Petri dish of interest, tiled into overlapping 416 × 416 pixel squares, and resulting tiles are fed serially to the trained plaque detection model. The detected plaques are deduplicated to account for the overlapping tiles which may have resulted in some plaques being detected twice, and the final annotated image is presented to the user. Optionally, the user may proceed with assay calculations within the application directly (for example, to determine phage titre) using the obtained plaque counts and dilution volumes. Figure created with BioRender. **(b)** Example image processed in the OnePetri mobile application. Three Petri dishes are detected with high confidence scores. Upon selecting a Petri dish, 155 phage plaques are enumerated and highlighted with a red box. Image provided by the Howard Hughes Medical Institute’s Science Education Alliance-Phage Hunters Advancing Genomics and Evolutionary Science (SEA-PHAGES) program.

### OnePetri rapidly and precisely enumerates bacteriophage plaques on a mobile device

In order to compare OnePetri’s accuracy to other currently available tools with a similar purpose, we benchmarked OnePetri against manual counts, as well as Plaque Size Tool, and CFU.AI, using a collection of 50 test images which the trained models would be seeing for the first time. One image had too many plaques to count manually (> 1,000) and was excluded from further analysis, despite OnePetri returning a value of 1,641 plaques, a seemingly accurate value.

OnePetri was benchmarked directly on-device and using the iOS development simulator included with the Xcode IDE on macOS. The time to result was significantly shorter when using OnePetri on-device versus Plaque Size Tool (*p*=0.0029), manual counts (*p*<0.001), and OnePetri in the iOS simulator (*p*=0.0041) (**Figure 3A**). No significant difference in time to result was seen when comparing OnePetri (on-device) to CFU.AI (*p*=0.4833). The mean time to result for OnePetri on-device was 1.91 seconds, OnePetri in simulator was 3.76 seconds, CFU.AI was 1.80 seconds, Plaque Size Tool was 3.21 seconds, and manual counts was 57.98 seconds (**Figure 3A**). OnePetri on-device was approximately 2x faster than the iOS development simulator, 1.7x faster than Plaque Size Tool, and 30x faster than manual counts, on average.

**Figure 3.**
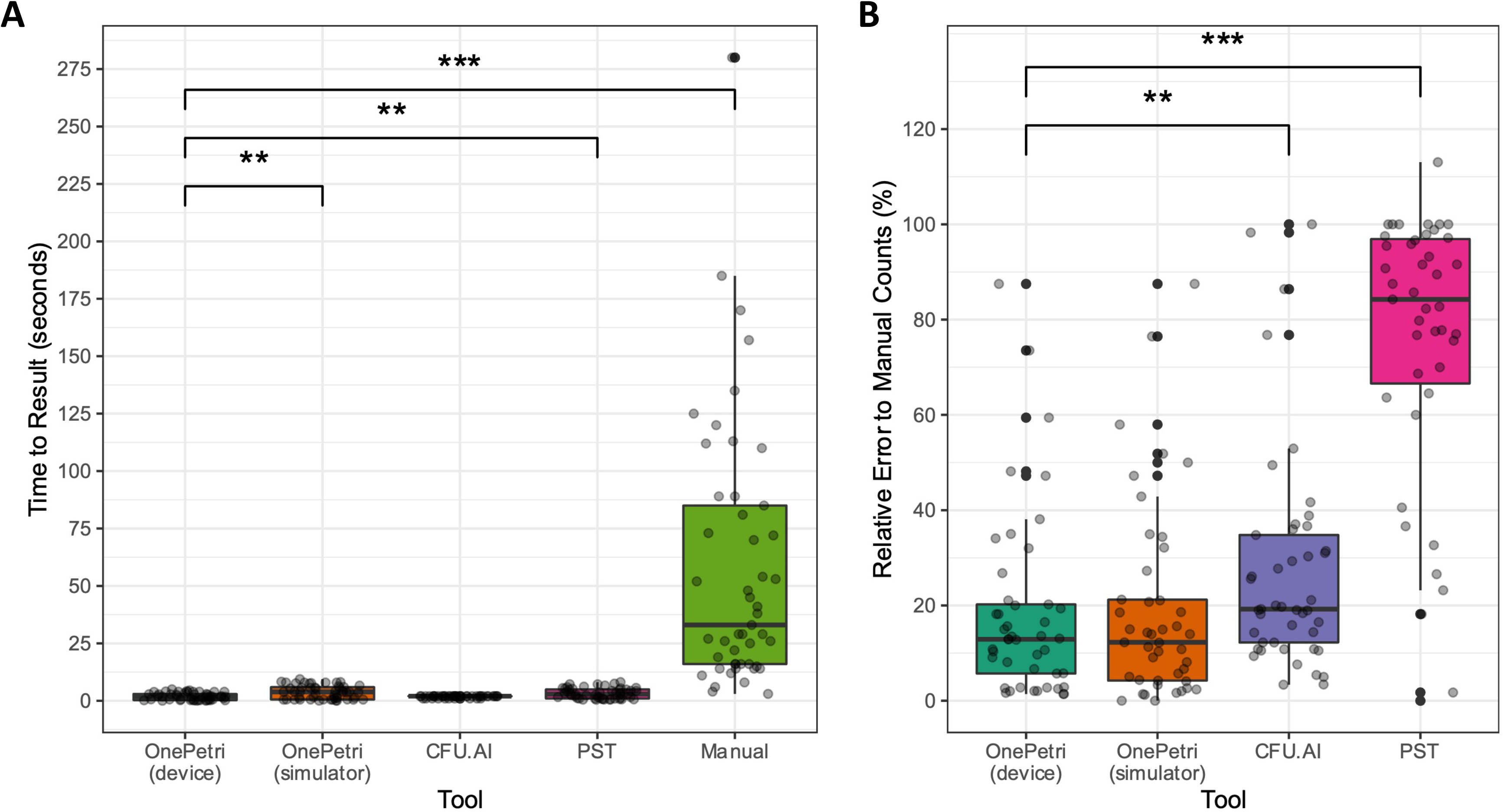
The OnePetri mobile application rapidly detects plaques with minimal error compared to other tools. OnePetri (on-device and in the iOS simulator) was benchmarked against CFU.AI, Plaque Size Tool (PST), and manual counts. The **(a)** total time to obtain plaque counts (in seconds), and **(b)** relative error rate of each tool (%), comparing counts from the tool to gold standard manual counts, were calculated and compared. (n = 49 images in the benchmarking dataset analyzed using each tool, * p ≤ 0.05, ** p ≤ 0.01, *** p ≤ 0.001, nonparametric one-way ANOVA using Wilcoxon rank sum test with Benjamini-Hochberg correction for multiple comparisons)

We compared the relative percent error of each approach to manual counts to get a sense of the overall accuracy of each tool. This value was calculated by taking the absolute value of the difference between actual and expected plaque counts, dividing by the expected plaque count, and multiplying by 100%. OnePetri on-device had the lowest median relative error of 12.90%, with a rate of 12.26% in the simulator, while CFU.AI and Plaque Size Tool had median relative errors of 19.23% and 85.71%, respectively, with these differences remaining significant after correcting for multiple comparisons (**Figure 3B**).

## Discussion

Phage enumeration through manual plaque counting has long been the gold standard in the field, despite the time-intensive nature of this approach. Over the years, several tools have been developed to help automate this approach, with varying levels of user intuitiveness and accuracy. Moreover, most tools have been developed for desktop computers, requiring Petri dish images to be uploaded to the computer for analysis, removing the user from their workflow in the laboratory. To this end, we developed OnePetri, a set of object detection models and a mobile application which can perform rapid plaque counting in a high-throughput fashion, directly in a laboratory environment, and which will improve regularly with ongoing model updates.

Using a diverse training dataset of Petri dish and plaque assay images from the HHMI SEA-PHAGES program, we were able to train YOLOv5s object detection models which could detect Petri dishes and phages plaques with high precision and recall (**Figure 1**). When benchmarked on a set of 50 images which the models have not been previously exposed to, OnePetri running on a physical iOS device was significantly faster for plaque counting than Plaque Size Tool and manual counting (**Figure 3A**). Using mean values for comparison, OnePetri on-device was approximately 30x faster than manual counts, representing significant time savings for the user, especially when analyzing multiple Petri dishes. Notably, OnePetri on-device was also significantly faster than using the iOS simulator included in the Xcode development suite, highlighting the need to benchmark iOS applications on-device rather than in simulators for accurate real-world values. No significant difference was seen in inference times between OnePetri on-device and CFU.AI. Despite having quicker detection times, OnePetri significantly outperformed CFU.AI and Plaque Size Tool in terms of relative error rates when comparing plaque counts from each tool to the true manual counts using the benchmarking image dataset (**Figure 3B**). The median error rate for OnePetri on-device (12.90%) was approximately 1.5x and 6.6x lower than CFU.AI (19.23%) and Plaque Size Tool (85.71%), respectively. No significant difference was observed between OnePetri error rates on-device versus in the iOS simulator. While a median error rate of 12.90% is quite low relative to the other tools, there remains room for improvement. Given the current error rate, we recommend that this version of OnePetri be used only when this level of error is acceptable for the assay at hand, and users should evaluate how OnePetri performs with their phage-host pairings before considering replacing manual counts entirely.

During benchmarking, we remarked that the Petri dish’s background surface (for example, on a dark lab bench, or held up against room light) can affect results, with more accurate counts being obtained when the Petri dish was against a dark surface. Furthermore, certain plaque morphologies, such as the “bull’s eye”, were sometimes incorrectly detected with the current version of OnePetri’s plaque detection model. OnePetri does require that images be of sufficient resolution for individual plaques to be distinguishable by the machine learning models, though we did not test this directly. However, all supported devices (modern smartphones from past 5-7 years) have camera resolutions which are well beyond what would likely be the minimum image size for reliable results.

The approach we used for image analysis on a mobile device allows users to serially analyze multiple Petri dishes within a single image, increasing throughput and reducing the time to results (**Figure 2**). Multiple detection parameters (object detection confidence thresholds and plaque deduplication overlap thresholds) can be easily changed within the application itself, allowing users to fine-tune the application’s performance to their unique setup, should the default values not be ideal.

The phage titration assay is the only assay currently supported within the mobile application. Upon entering the volume of sample plated and corresponding plate dilutions, the initial phage titre is calculated based on the number of plaques present on serially diluted plates. Support for additional phage and bacterial assays is planned for late-2021/early-2022. OnePetri does not currently support spot assays for approximating phage titre and requires that each Petri dish contain phage of a single dilution, as all plaques on a given dish are counted assuming they are from the same diluted sample. Unlike Plaque Size Tool, OnePetri does not currently directly measure or infer individual plaque size. This may be added in a future version of OnePetri, along with support for exporting a summary report of all Petri dishes analyzed in a given session. A version of the OnePetri mobile application which supports Android devices is currently under development and should be released early-2022.

## Conclusion

We present a pair of trained object detection machine learning models for the identification of Petri dishes and phage plaques, as well as OnePetri, a mobile application for iOS which leverages these models for the rapid and reproducible enumeration of phage plaques. OnePetri is now freely available to download from the Apple App Store on iOS. The application source code, trained models, training data, and benchmarking dataset and analysis scripts are all available for download under open-source licenses. When compared to the manual counting gold standard, as well as CFU.AI and Plaque Size Tool, OnePetri had minimal relative error with significantly lower time to results.

## Acknowledgements

This work was supported by the Natural Sciences and Engineering Research Council of Canada (NSERC), the Fonds de recherche du Québec – Nature et technologies (FRQNT; #282402), and AbbVie Canada to M.S. We are especially grateful to the Howard Hughes Medical Institute’s Science Education Alliance-Phage Hunters Advancing Genomics and Evolutionary Science (SEA-PHAGES) program for proving the large collection of Petri dish and plaque assay images to be used for model training and testing—this project would not have been possible without their generous contribution. We would also like to thank Hendrik Kueck (Pocket Pixels Inc.) for his guidance on topics in computer vision and machine learning object detection model training, as well as Mohamed Traore (Roboflow Inc.) and the entire Roboflow team for their support and for providing access to their image annotation and preprocessing platform. We would like to acknowledge those who helped test and provided feedback on OnePetri for iOS during the beta testing period, prior to its official release.

## References

1. Kropinski AM, Mazzocco A, Waddell TE, Lingohr E, Johnson RP. Enumeration of Bacteriophages by Double Agar Overlay Plaque Assay. In: Clokie MRJ, Kropinski AM (eds). Bacteriophages: Methods and Protocols, Volume 1: Isolation, Characterization, and Interactions. 2009. Humana Press, Totowa, NJ, pp 69–76.

2. Abedon ST, Yin J. Bacteriophage Plaques: Theory and Analysis. In: Clokie MRJ, Kropinski AM (eds). Bacteriophages: Methods and Protocols, Volume 1: Isolation, Characterization, and Interactions. 2009. Humana Press, Totowa, NJ, pp 161–174.

3. Culley S, Towers GJ, Selwood DL, Henriques R, Grove J. Infection Counter: Automated Quantification of in Vitro Virus Replication by Fluorescence Microscopy. Viruses 2016; 8.

4. Katzelnick LC, Coello Escoto A, McElvany BD, Chávez C, Salje H, Luo W, et al. Viridot: An automated virus plaque (immunofocus) counter for the measurement of serological neutralizing responses with application to dengue virus. PLoS Negl Trop Dis 2018; 12: e0006862.

5. Trofimova E, Jaschke PR. Plaque Size Tool: An automated plaque analysis tool for simplifying and standardising bacteriophage plaque morphology measurements. Virology 2021; 561: 1–5.

6. Cacciabue M, Currá A, Gismondi MI. ViralPlaque: a Fiji macro for automated assessment of viral plaque statistics. PeerJ 2019; 7: e7729.

7. Russell DA, Hatfull GF. PhagesDB: the actinobacteriophage database. Bioinformatics 2017; 33: 784–786.

8. Paszke A, Gross S, Massa F, Lerer A, Bradbury J, Chanan G, et al. PyTorch: An Imperative Style, High-Performance Deep Learning Library. In: Wallach H, Larochelle H, Beygelzimer A, Alché-Buc F d’, Fox E, Garnett R (eds). Adv. Neural Inf. Process. Syst. 2019. Curran Associates, Inc.

9. Jocher G, Stoken A, Borovec J, NanoCode012, Chaurasia A, TaoXie, et al. ultralytics/yolov5: v5.0 - YOLOv5-P6 1280 models, AWS, Supervise.ly and YouTube integrations. 2021. Zenodo.

10. H. Rezatofighi, N. Tsoi, J. Gwak, A. Sadeghian, I. Reid, S. Savarese. Generalized Intersection Over Union: A Metric and a Loss for Bounding Box Regression. 2019 IEEECVF Conf. Comput. Vis. Pattern Recognit. CVPR. 2019. pp 658–666.

